# Functional evolutionary convergence of long noncoding RNAs involved in embryonic development

**DOI:** 10.1101/2022.06.15.496228

**Authors:** Ane Olazagoitia-Garmendia, Rodrigo Senovilla-Ganzo, Fernando Garcia-Moreno, Ainara Castellanos-Rubio

**Affiliations:** University of the Basque Country, UPV-EHU, Leioa, Spain; Biocruces Health Research Institute, Barakaldo, Spain; Achucarro, Basque Center for Neuroscience, Leioa, Spain; Ikerbasque, Basque Foundation for Science, Bilbao, Spain; CIBERDEM, Madrid, Spain

## Abstract

Long noncoding RNAs (lncRNAs) have been identified in almost all vertebrates, but the functional characterization of these RNA molecules is being challenging, mainly due to the lack of linear sequence homology between species. In this work, we aimed to find functional evolutionary convergent lncRNAs involved in development by screening of k-mer content (non linear similarity) and secondary structure-based approaches combined with *in silico, in vitro* and *in vivo* validation analysis. From the currently identified Madagascar gecko genes, we found a lncRNA with a similar k-mer content and structurally concordant with the human lncRNA *EVX1AS*. Analysis of function related characteristics together with locus-specific targeting of human and gecko *EVX1AS* (i.e. CRISPR Display) in human neuroepithelial cells and chicken mesencephalon confirmed that gecko *Evx1as-like* lncRNA mimics human *EVX1AS* function and induces *EVX1* expression independently of the target species. Our data show functional conservation of non-homologous lncRNAs and presents a useful approach for the definition and manipulation of lncRNA function within different model organisms.

## Introduction

Long noncoding RNAs (lncRNAs) are RNA molecules longer than 200bp in length that do not have coding potential (Rinn and Chang, 2012). LncRNAs are gaining importance due to their involvement in a wide range of biological processes, and some of them have been described to be implicated in different aspects of embryonic development (Fico et al., 2019). However, the study of lncRNA relevance through their evolutionary conservation has been challenging due to their lack of linear sequence homology among species. Evolutionary conservation is widely used as an indicator of the functional significance of newly discovered genes, and the simple search for homology at the nucleotide level has proven to be valuable for protein-coding genes. However, lncRNAs with similar functions often lack linear sequence homology which implies lncRNA function cannot be readily assigned from their nucleotide sequence. K-mer based comparison methods have been demonstrated to be useful to find functionally related lncRNAs with different spatial arrangements of related sequence motifs, where a k-mer is defined as all possible combinations of a continuous sequence of nucleotides of a given length k. K-mer-based classification has been demonstrated to be a powerful approach to detect recurrent relationships between motif sequence and function in lncRNAs even in the lack of evolutionary conservation (Kirk et al., 2018; Sprague et al., 2019). Additionally, lncRNAs with the same secondary or tertiary structure can exert identical molecular functions despite divergent nucleotide sequences (Smith et al., 2013), thus analyzing structural equivalence could also help identify lncRNAs with evolutionary preserved mechanisms (Ponti et al., 2018). Moreover, the molecular role of lncRNAs is tied to other characteristics such as the subcellular localization, the abundance within the cell or the interactions with other molecules (Much et al., 2022; Statello et al., 2021).

In this work, we have taken advantage of k-mer and structure similarity analyses to find evolutionary convergent lncRNAs involved in development. For this purpose, we used Madagascar ground gecko (*Paroedura pictus*) as the target species to find functionally conserved lncRNAs related to embryonic development. The Madagascar gecko is a useful species to investigate the evolutionary path of vertebrate features due to its phylogenetic position within the *squamates* order of reptiles (Nomura et al., 2013; Noro et al., 2009). Additionally, we have performed several *in vitro* and *in vivo* analyses to investigate the functional cross-species conservation of the candidate lncRNAs. For the *in vivo* analyses, we have used chicken (*Gallus gallus*) embryos as an extra-phyletic species to both human and gecko species. The chicken is the most suitable animal model for experimental embryology, and it is a perfect playground for genetic engineering during development (García-Moreno et al., 2018; Itasaki et al., 1999; Nakamura and Funahashi, 2013). Using these approaches, we found that the lncRNA *EVX1AS* is evolutionary convergent between species, and it regulates its divergent coding gene *EVX1* independently of the origin of the lncRNA and the recipient species. This evolutionary convergence implies that these two lncRNAs emerged independently in the two species, evolving in parallel to play equivalent functions from non-homologous sequences.

## Results

### Nonlinear sequence similarity between human and gecko embryogenesis lncRNAs

We hypothesized that functionally related lncRNAs involved in embryonic development could harbor related motif contents, although lacking linear sequence similarity. To test this, we used SEEKR (sequence evaluation from k-mer representation) webapp (Kirk et al., 2018) to find development associated lncRNAs (with already described functions in humans) that could be functionally convergent between human and gecko. Using the All human lncRNA (Gencode v41) set as a normalization set, we calculated the k-mer profile at k=3 and k=4 k-mer lengths of human *NEAT1, MEG3* and *EVX1AS* against all sequences from *P.picta* genome v1 in order to find equivalent lncRNAs (Hara et al., 2018). For *MEG3* and *EVX1AS*, the genes with the highest k-mer values in both k-3 and k-4 comparisons (above 0.65 similarity for k-3 and above 0.5 similarity for k-4) were chosen, and the derived isoforms were evaluated for their noncoding nature using the Coding Potential Calculator (Kang et al., 2017). For *MEG3* and *EVX1AS*, candidate noncoding transcripts were selected based on their noncoding nature and its score in both k=3 and k=4 analyses (*Pp-Meg3* and *Pp-Evx1as* from now on). Both candidates were present in the highest percentiles (>99%ile) with k-mer values of 0.66 and 0.81 for k-3 and 0.52 and 0.61 for k-4 respectively (Fig. 1A). However, no such candidate was found for *NEAT1*.To verify the validity of this approach, we also compared the k-mer content distribution between human and mouse lncRNAs as they are known to be functionally conserved. These comparisons resulted in comparable percentile values to those found between human and gecko lncRNAs with 100 %ile for *MEG3* and >97%ile for *EVX1AS*. To evaluate the specificity of our results, we also performed the reverse analysis and compared our gecko candidates with the human set of all human lncRNAs. The results of this reversal SEEKR analysis, showed that our lncRNA candidates maintain a high k-mer correlation (higher than 0.8 and 0.55 for both k-3 and k-4 respectively) and they are in the highest percentile (>97) (Sup. Fig. 1A). All together, these results suggested these ^2^selected gecko noncoding RNAs are the most likely functional equivalents to human lncRNAs and ruled out the possibility of the candidates having been selected by chance.

**Figure 1:**
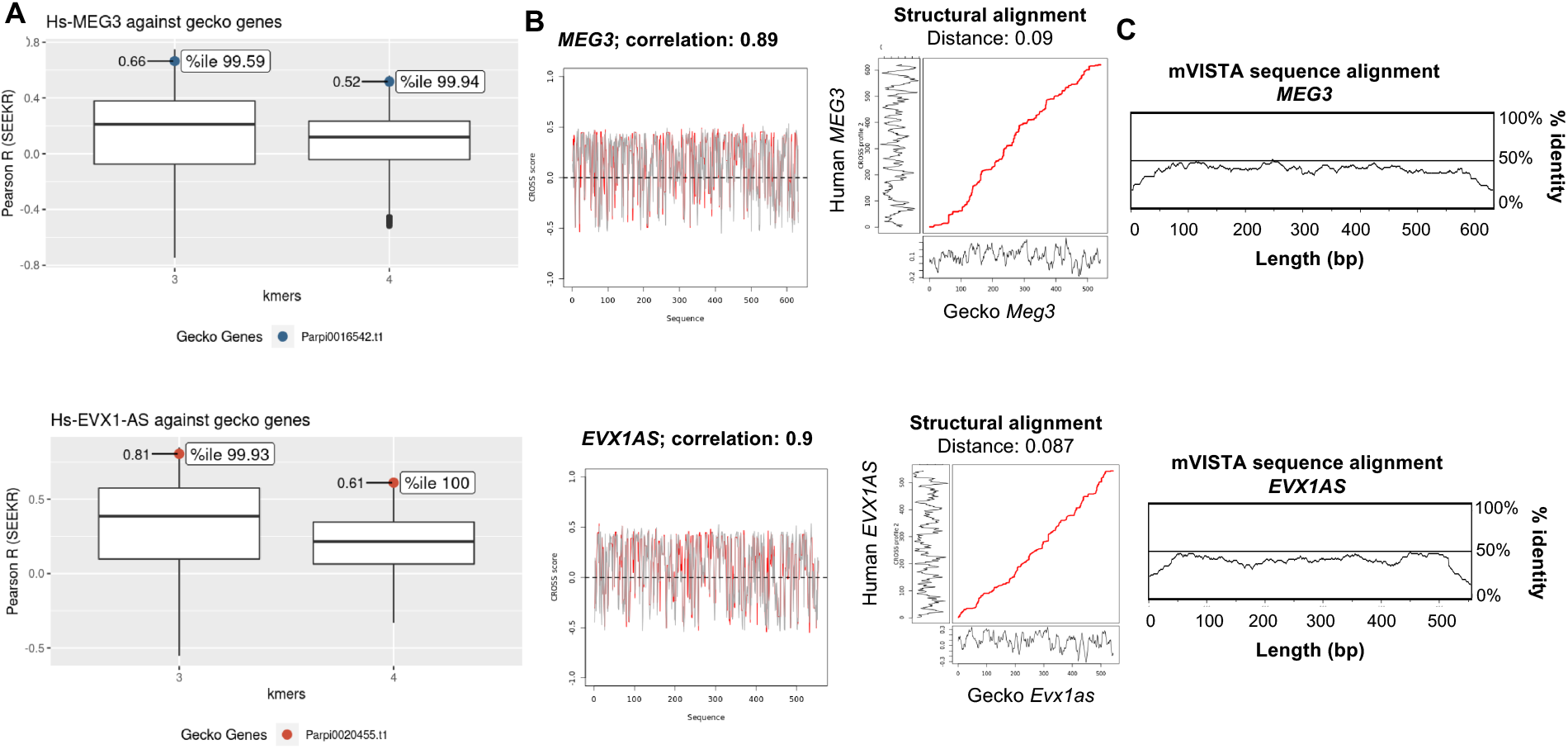
Nonlinear sequence similarity between human and gecko embryogenesis lncRNAs. **(A)** k=3 and k=4 k-mer content analysis of *MEG3* and *EVX1AS* lncRNAs using SEEKR tool. Candidate lncRNAs were compared against the Madagascar gecko genome. The boxplot represents all gecko genes, and the dots highlight the position of the selected candidate genes as well as its correlation values (Pearson, SEEKR) and its percentile (%ile). Blue and red dots correspond to the *MEG3* and *EVX1AS* genes respectively. **(B)** Secondary structure similarity study using CROSSALIGN tool. Optimal matching region by OBE-DTW algorithm (left) and overall structural similarity by Standard-DTW (right). Structural profiles are obtained with CROSS Global Score for the two RNAs (score >0 means a double-stranded nucleotide; <0 single-stranded). **(C)** Linear conservation analysis of *MEG3* (top) and *EVX1AS* (bottom) performed using mVISTA. Human sequence is shown on the x-axis and percentage similarity to the corresponding gecko sequence on the y-axis. The graphical plot is based on sliding-window analysis of the underlying genomic alignment. A 100-bp sliding window is at 25-bp nucleotide increment is used.

Given that the gecko noncoding transcripts were 6-to-8 times longer than their human counterparts (4448 bp vs 554bp for *EVX1AS* and 3751 bp vs 632bp for *MEG3)*, we used CROSSalign (Ponti et al., 2018) to identify regions of structural similarity between the different length profiles (Fig. 1B). Both lncRNA pairs presented a structural distance lower than 0.095 (where 0 means identical structural profiles) and a 90% structure correlation. The normalized structural distance between the secondary structure profiles of *EVX1AS* was calculated as 0.087 (p-value = 0.001) with a correlation of 90%. In the case of *MEG3* the distance was 0.09 (p-value=0.01) with a correlation of 89%. Dinucleotide shuffled sequences of the gecko lncRNAs showed structural distances higher than 0.09 in both cases. Interestingly, we did not find any specific regions of conventional linear sequence homology between the pairs of structurally equivalent lncRNAs by mVISTA using the 70% conservation over a 100bp window criteria, neither by a less stringent DotPlot analysis (with a 20% similarity) (Fig. 1C and Sup. Fig. 1B) (Frazer et al., 2004; Rice et al., 2000); suggesting an independent evolutionary origin of both transcripts. Moreover, analysis of the genomic neighborhood of the gecko lncRNA confirmed that the genomic context in which it is located is different from that of human *EVX1AS* (non-common synteny), corroborating the lack of homology of these lncRNAs. (Sup. Fig. 1C).

Thus, our analysis revealed the existence of disparately originated pairs of lncRNAs in varied vertebrate species. These two lncRNAs were predicted to act upon pairs of orthologous genes and presented very significant levels of structural similarity and k-mer content. Together, it suggested that these lncRNAs could play similar functions despite the lack of evolutionary relationship.

### Human and gecko EVX1AS are principally expressed in brain

In order to verify if the gecko lncRNA candidates were actually expressed, we quantified the expression of the *Pp-Meg3* and *Pp-Evx1as* in different tissues from pre-hatching geckos. *Pp-Meg3* was only expressed in the tail and the carcase (Fig. 2A), with a very similar expression level in the two tissues. Expression analysis of *Pp-Evx1as* demonstrated that this lncRNA is widely and tissue-specifically expressed in this species, showing the highest expression in the brain, 9 to 280 times higher than in the rest of the tissues (Fig. 2A, Sup. Fig. 2A). In mice, lncRNA *Evx1as* has been described to transcriptionally regulate its nearby coding gene *Evx1* (Luo et al., 2016). Analysis of gecko *Evx1* expression in the embryonic tissues showed the wide characteristic expression of this coding gene, with the highest expression levels in the heart and the carcase and the lowest in the lung (Fig. 2A, Sup. Fig. 2A). We also analyzed the expression of both *EVX1AS* and *EVX1* in an RNA pool of different human tissues purchased from Clontech. As previously observed in gecko embryos, *EVX1AS* was widely expressed among the different tissues with the highest expression also present in the brain (with 1.5-12 times higher values) and the lowest in the colon and the thymus (Fig. 2B, Sup. Fig. 2B). Regarding the coding gene *EVX1*, it likewise presented a broad tissue expression with the highest levels in the kidney and the lowest also in the lung (Fig. 2B, Sup. Fig. 2B). Given that human brain tissue is not an accessible material, we also evaluated the expression of *EVX1AS* and *EVX1* in the human neuroepithelial cell line SHSY5Y. Both genes were expressed in this cell line confirming the validity of this culture model for further *EVX1AS* analysis (Fig. 2C).

**Figure 2:**
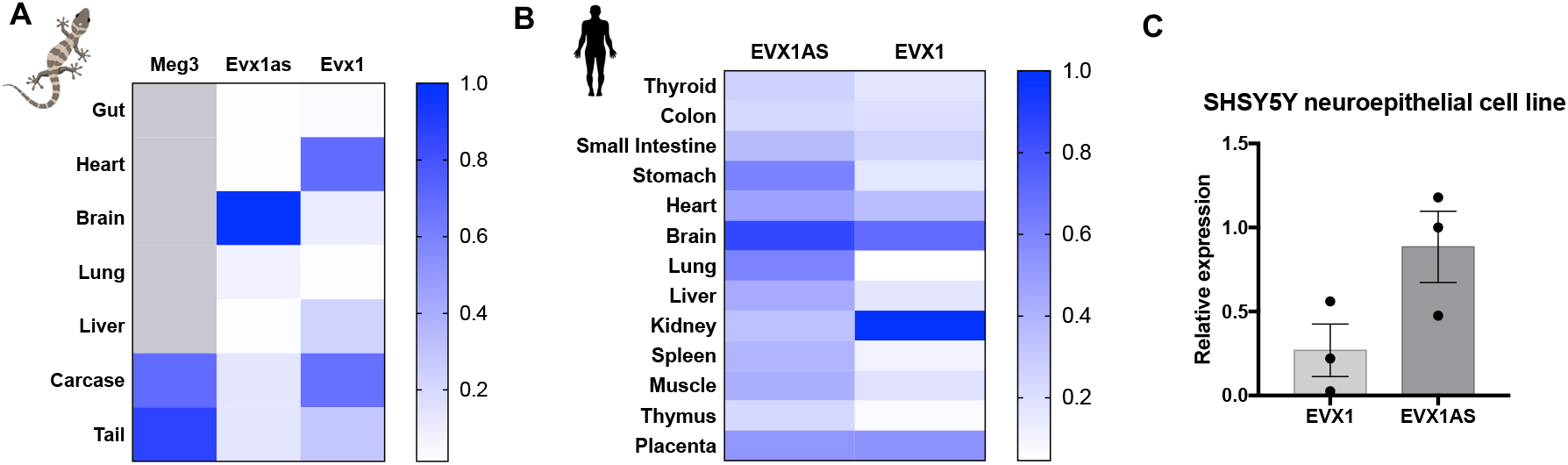
Human and gecko EVX1AS are principally expressed in the brain. Heat map showing relative expression (from 0 to 1) of **(A)** gecko *Meg3, Evx1as* and *Evx1* and **(B)** human *EVX1AS* and *EVX1* in a range of tissues. **(C)** *EVX1AS* and *EVX1* expression analysis in the human neuroepithelial cell line SHSY5Y. Data represents the media and standard error of 3 independent analyses.

Given the multi-level similarities, from motif content and structure to tissue expression pattern, we selected *Pp-Evx1as* lncRNA for further analysis of its potentially shared function. LncRNAs showing common function -among other shared features-but lacking linear sequence homology would indicate these lncRNAs evolved by functional convergence: they evolved separately, from independent ancestor sequences, but converged into an equivalent function in parallel.

### Human and gecko EVX1AS share function-related characteristics

RNA structure, subcellular localization and abundance of lncRNAs are generally related to their function and molecular roles. Thus, to experimentally assess their functional convergence, we evaluated several function-related characteristics (structure, localization and abundance) for both human and gecko *EVX1AS*. These two lncRNA forms showed an almost identical migration pattern when *in vitro* transcribed lncRNAs corresponding to the structural analogue region were migrated in a non-denaturing agarose gel (Fig. 3A). Conversely, an unrelated in vitro transcribed lncRNA, which was used as a control, showed a totally different migration pattern, confirming the equivalency of the secondary structures of human and gecko *EVX1AS* predicted in silico (Fig. 1B). For subcellular localization assessment, the amounts of *EVX1AS* were quantified in whole and nuclear fractions of human neuroepithelial cells and the gecko brain. As previously described (Luo et al., 2016), *EVX1AS* was found in both cellular compartments suggesting that apart from regulating *EVX1* in cis, this lncRNA also exerts other yet undescribed biological functions. The percentage of *EVX1AS* present in the nucleus was very similar in both species (approximately 30%) (Fig. 3B), further supporting its functional convergence. Evaluation of *EVX1AS* abundance was performed using human and gecko brain cDNA and a reference plasmid of each lncRNA (Sup. Fig. 3A). We had the limitation that while the gecko brains tested were embryonic, human brain samples were from adult individuals. However, the amount of *EVX1AS* molecules per cell in both cases was lower than 1 (Fig. 3C), pointing to a cell type specific expression of the lncRNA within the brain. The higher amount of copies per cell present in the gecko brain may be suggestive of a high peak of activity of this lncRNA regulating *EVX1* during embryonic development.

**Figure 3:**
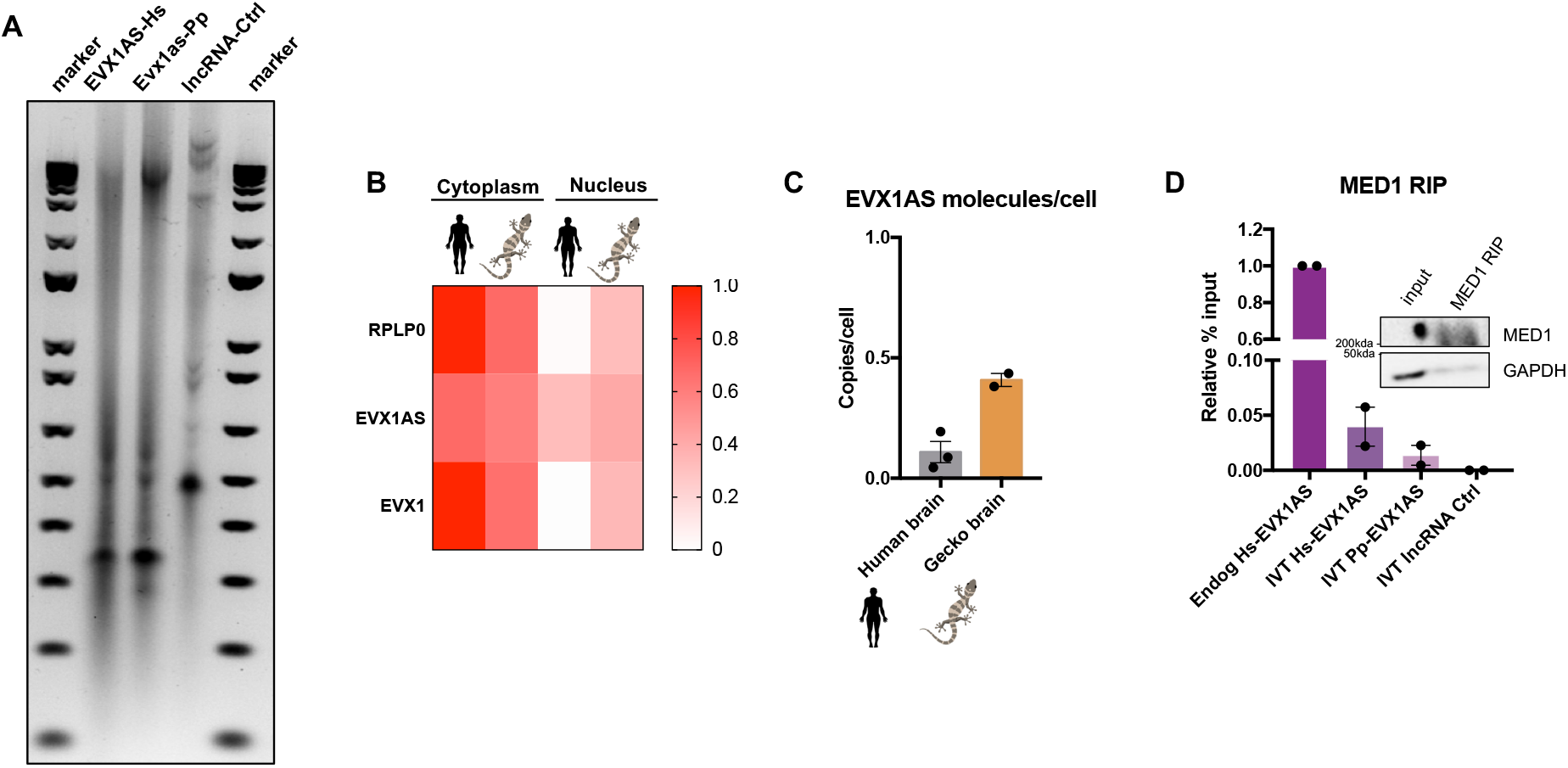
Human and gecko EVX1AS share function-related characteristics. **(A)** Mobility assay of in vitro transcribed (IVT) human (Hs) and gecko (Pp) *EVX1AS*. A non related lncRNA was used as a control (Ctrl). **(B)** Cellular localization of human and gecko *EVX1AS* in human SHSY5Y cells (n=3) and gecko brains (n=2). *PO* and *EVX1* were used as cytoplasmic controls. Analysis of *EVX1AS* copy number in human and gecko brain. Data represents the mean and standard error of the experiments. **(D)** Analysis of endogenous and in vitro transcribed *EVX1AS* levels bound to MED1 protein after immunoprecipitation. An in vitro transcribed unrelated lncRNA was used as a control. A representative western blot of the immunoprecipitation is shown.

The regulation of mouse *Evx1* expression by *Evx1as* has been described to be mediated by the binding of the lncRNA to the mediator complex (Luo et al., 2016). The mediator complex is a multiprotein complex that functions as a transcriptional coactivator and interacts with a wide range of proteins. MEME motif analysis of our lncRNAs, showed enrichment of several brain function related protein binding motifs, as MATR3, HNRNPA1 and PTBP1 (Sup. Fig. 3B), that have been reported to interact with members of the mediator complex according to GENEMANIA server (Warde-Farley et al., 2010). Thus, to analyze if both our lncRNA forms also bind the mediator complex, we performed an RNA immunoprecipitation experiment of MED1 protein using lysates from SHSY5Y cells. Immunoprecipitation of human MED1 was able to retrieve both, endogenous and IVT human *EVX1AS*. Additionally, we also observed the lncRNA-MED1 interaction when IVT *Pp-Evx1as* was added to the protein lysates. Conversely, our IVT lncRNA control did not interact with MED1, confirming the specificity of the binding (Fig. 3D).

Equivalence in structure, subcellular localization, and abundance of *EVX1AS* from both gecko and human, together with the interaction with the mediator complex, strongly suggested a shared cellular function.

### EVX1AS is functionally convergent among species

To analyze whether the previous results were representative of a functional in-cell equivalence, we conducted overexpression experiments of both lncRNAs in two experimental paradigms: *in vitro*, in SHSY5Y cells; and *in vivo*, in chick embryonic brains. For this approach, we used the CRISPR-Display technique (Shechner et al., 2015), in order to specifically localize *EVX1AS* into the promoter of human or chicken *EVX1*. For *in vitro* experiments, we cloned either the human or gecko *EVX1AS* lncRNA preceded by two different guide RNAs specific for human *EVX1* promoter (HS1 and HS2 being sgRNAs targeting human *EVX1* TSS1 and TSS2 respectively). Otherwise, for *in vivo* experiments, gRNAs for chicken *Evx1* promoter were designed (GG1 and GG2 as sgRNAs targeting chicken *Evx1* TSS1 and TSS2 respectively). These constructs were followed by a 3’ box and cloned into a CMV driven vector to guide and tether the *EVX1AS* RNA to the *EVX1* promoter (Sup. Figure 4A, B). The transfection of Hs-*EVX1AS* vectors into the SHSY5Y cells showed a 4-13×10^6^ fold statistically significant overexpression (p<0.01 for HS1 and p<0.001 for HS2) (Fig. 4A). Using Pp-*Evx1as* plasmids, 1.5-3×10^6^ fold overexpression could be achieved, being significant only with HS2 gRNA (p<0.001) (Fig. 4B). The directed overexpression of the human *EVX1AS* was able to induce the expression of *EVX1* mRNA in the SHSY5Y human neuroblastoma cell line about 2 times (p<0.01 for HS2) (Fig. 4A). When gecko *Evx1as* (*Pp-Evx1as*) was overexpressed in the human cells, we also observed a 1.5-2-fold increase in the mRNA levels of *EVX1* (p<0.01 for HS2) (Fig. 4B), similar to what it was previously observed using this same approach (Luo et al., 2016). No induction of *GAPDH* negative control could be observed in neither of the cases (Sup. Fig. 4E,F) confirming that these two lncRNAs functionally converged in their role of controlling *EVX1* mRNA expression. Interestingly, the tethering mediated by the sgRNA2 (named HS2) (closer to the *EVX1* start codon) performed more efficiently, independently of the origin of the lncRNA.

**Figure 4:**
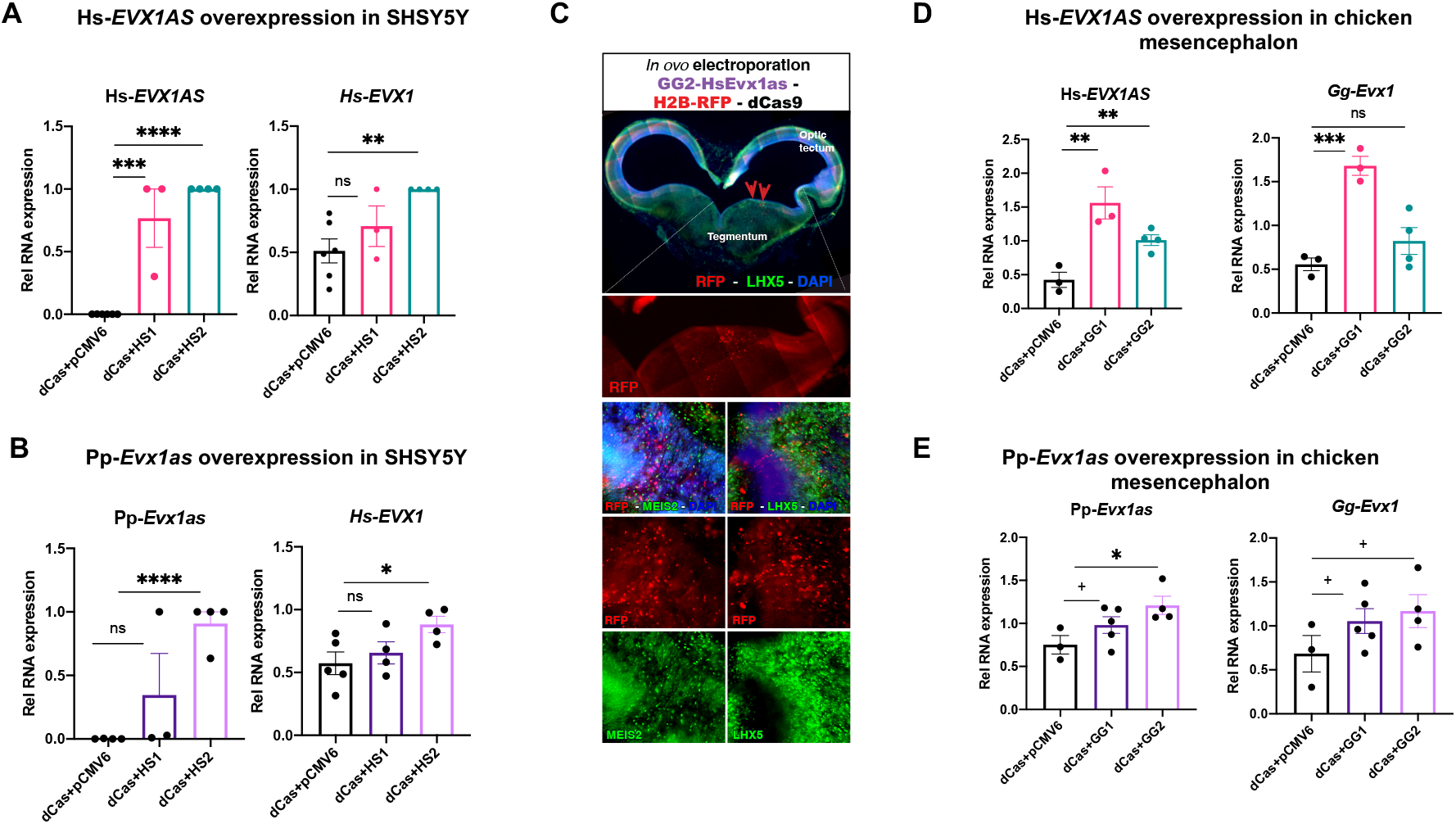
EVX1AS is functionally convergent among species. **(A)** Relative RNA expression of human *EVX1AS* lncRNA and *EVX1* upon human *EVX1AS* overexpression in SHSY5Y cells. **(B)** Relative RNA expression of gecko *Evx1as* lncRNA and human *EVX1* upon gecko *Evx1as* overexpression in SHSY5Y cells. **(C)** Immunohistochemistry of the chicken embryonic midbrain on coronal section showing the successful transfection of the lncRNA plasmid. Transfected cells are shown in red due to their ectopic expression of red fluorescent protein RFP. LHX5 and MEIS2 patterns of expression (in green) indicate the mesencephalic region that was actually transfected. DAPI counterstain in blue. **(D)** Relative RNA expression of human *EVX1AS* lncRNA and chicken *Evx1* upon human *EVX1AS* overexpression in chicken mesencephalon. **(E)** Relative RNA expression of gecko *Evx1as* lncRNA and chicken *Evx1* upon gecko *Evx1as* overexpression in chicken mesencephalon. Data represents the mean and standard error of 3-6 independent experiments. (+p<0.1, *p<0.05, **p<0.01, ***p<0.001, ****p<0.0001, ns: non-significant according to a Student’s t-test). HS1 and HS2: sgRNAs targeting human *EVX1* TSS1 and TSS2 respectively. GG1 and GG2: sgRNAs targeting chicken *Evx1* TSS1 and TSS2 respectively.

To further confirm these data, we decided to use an extra-phyletic *in vivo* model that could act as a recipient for both species. The chicken embryo is a good model for these analyses as it represents a suitable experimental model which is phylogenetically distant from both human and gecko. As it has been previously described we found expression of the *Evx1* coding gene in the chicken embryo mesencephalon, but not in the forebrain (Sup. Fig 4C). We electroporated our lncRNAs together with the catalytically dead Cas9 into the mesencephalon of embryonic day 3 (E3) chicken embryos, when the mesencephalic *Evx1* expressing cells are mostly being generated (Agarwala et al., 2001; Fogel et al., 2008). We validated our strategy using immunofluorescence, as we confirmed that we were able to electroporate and express our plasmids in the mesencephalic region of chicken embryos as determined by LHX5 and MEIS2 expression (Fig. 4C and Sup. Fig. 4D). Three days after overexpression (E6), as previously observed in the human cells, the tethering of both human and gecko *EVX1AS* to the chicken *Evx1* promoter induced the levels of *Evx1* mRNA between 1.2 and 3.5 fold (p<0.001 for Hs-EVX1AS and p<0.08 for Pp-Evx1as) (Fig. 4D,E), while *Gapdh* expression did not vary significantly (Sup. Fig. 4G,H); confirming the functional convergence of the human and gecko lncRNAs.

The ability of the human and gecko *EVX1AS* to regulate the expression of human and chicken orthologous coding gene *EVX1* in all the combinations (human-human, gecko-human, human-chicken, gecko-chicken) confirms the functional similarity of both lncRNAs, and points to a case of evolutionary convergence.

## Discussion

LncRNAs regulate fundamental cellular processes, such as embryonic development, but still the physiological importance of the majority of lncRNAs remains to be identified. The lack of strategies to find relationships between lncRNA sequence and mechanism makes it difficult to functionally classify them. Additionally, the mechanistic studies using human material have also great limitations, making the functional characterization of lncRNAs a very difficult task. Moreover, the absence of linear conservation for these transcripts makes the identification of ortholog lncRNAs among different species very challenging, especially between evolutionarily distant species such as gecko and human. Thus, the need to develop a strategy to investigate functional convergent lncRNAs, bearing in mind that lack of sequence conservation does not directly imply different functions, will be of great importance to fully understand the evolution and function of these emerging regulatory elements.

In this study, we used SEEKR algorithm for nonlinear comparison of lncRNAs to identify potential gecko functional counterparts of human lncRNAs involved in development. Additionally, we used CROSSAlign to confirm the structural similarity among the selected candidates. Collectively, our data support that lncRNAs from different species can perform similar functions through vastly different linear sequences, and that nonlinear sequence comparisons can be used to discover evolutionary convergent lncRNAs among species. Moreover, the information about lncRNA structure and the identification of structurally equivalent regions has been useful to complement sequence analysis and to translate *in silico* analysis to *in vitro* and *in vivo* experiments.

Nonlinear sequence similarity and structure analyses of the three analyzed embryonic development-related human lncRNAs resulted in two candidate gecko lncRNAs that could perform the same functions as human lncRNAs *MEG3* and *EVX1AS. MEG3* has been involved in pluripotency and reprogramming and it is mainly expressed in the hypophysis (Stadtfeld et al., 2010; Zhou et al., 2010). The expression analysis of *Pp-Meg3* in the gecko tissues only showed expression in the tail and the carcase suggesting that we were not looking at a functional homolog. *EVX1AS* has been related with the mesoderm differentiation and it is known to regulate the expression of *EVX1* coding gene, which is expressed in the midbrain during embryonic development (Bell et al., 2016; Luo et al., 2016). We found that *Pp-Evx1as* was mainly expressed in the embryonic gecko brain, being a suitable candidate for further function related studies.

As previously stated, lncRNAs acquire complex structures that usually dictate their function. Moreover, lncRNAs can be found throughout the cell, acting in a wide range of cellular processes, and are commonly lowly expressed (Cabili et al., 2015; Much et al., 2022). These characteristics-structure, subcellular location, level of expression-would be common to those lncRNAs that exert the same functions, helping in the evaluation of putative convergent lncRNAs. The identical gel migration pattern of *Pp-Evx1as* and *Hs-EVX1AS*, together with the ubiquitous localization and the scarce expression of both lncRNAs suggested that they could be playing parallel functions. Moreover, these lncRNAs presented common motifs known to recruit RNA binding proteins, providing additional insight into similarities between the two lncRNAs. Interestingly, the three RNA-binding proteins predicted to bind the human and gecko lncRNAs showed interactions with different members of the mediator complex, as reported by the GENEMANIA server (Warde-Farley et al., 2010). Consistent with these data, we were able to show that human and gecko *EVX1AS* physically interact with MED1 protein, which has been described to bind *EVX1AS* for *EVX1* transcription regulation (Luo et al., 2016), further supporting the idea of their functional similarity.

*EVX1AS* is known to be involved in regulatory processes within the nucleus, i.e. regulating the expression of its nearby coding gene *EVX1*. Although the mechanistic dissection of these types of regulators *in vivo* is technically challenging, we took advantage of the CRISPR-Display technique, which is a very useful tool for relocating lncRNA transcripts to an ectopic site (Shechner et al., 2015). This technique allowed us to direct *EVX1AS* from different species (human and gecko) to the promoter of human and chicken *EVX1* coding gene, regardless of the lncRNA and recipient species. Using this approach, we showed that *Pp-* and *Hs-EVX1AS* are able to regulate *EVX1* expression in both model systems, human and chicken, supporting their functional similarity. Remarkably, these lncRNA are not derived from a common lncRNA ancestor, as evidenced by their lack of sequence homology. Instead, they evolved independently, in parallel, and their structures and functions converged in the different species tested.

Overall, this study provides insight into the functional similarity of embryonic development related lncRNA *EVX1AS* by analyzing its k-mer content and secondary structure together with the study of its expression, localization and abundance in human and gecko. Additionally, *in vitro* and *in vivo* studies confirmed the evolutionary convergence of these lncRNAs. The existence of non-homologous lncRNAs that share a common function and other features strongly suggest that evolutionary convergence played a role in the diversification of lncRNAs.

This approach can be of great use to describe the functional properties of lncRNA molecules within different species, facilitating the study of a wide range of biological processes, such as embryonic development, in diverse model organisms. To our knowledge, this is the first time that functional convergence of non-homologous lncRNAs has been shown and initiates the search for equivalent RNA evolution variants.

## Material and methods

### Selection of candidates lncRNAs using k-mer content

Starting from previously described embryonic development-related lncRNAs, we decided to analyze *NEAT1, MEG3* and *EVX1AS* based on their length (<4kb) and their already described conservation between mouse and human. We then analyzed the k-mer content of our candidate lncRNA sequences against all the sequences of our reptilian model, the Madagascar ground gecko (*Paroedura pictus*) genome v1 (Hara et al., 2018), using online available webapp SEEKR interface. We used all human lncRNAs from GENCODE as a background set to derive the mean and standard deviation of the counts for each k-mer (Kirk et al., 2018). Additionally, we did our analysis using two k-mer lengths: k=3 and k=4. The gecko genes with highest k-mer values in both k-mer lengths, k=3 and k=4, were chosen (0.66 and 0.81 for k-3 and 0.52 and 0.61 for k-4 respectively) (Fig. 1A) and the noncoding nature of the derived gecko transcripts was analyzed using the Coding Potential Calculator (CPC) web-based interface (Kang et al., 2017). The reverse comparison, in which we analyzed our gecko lncRNAs of interest against all human lncRNAs and the k-mer content correlations between the mouse and human lncRNA forms were performed as controls. All boxplots from SEEKR results displayed in the results and supplementary figures were created in *R*, using *ggplot2* package and SEEKR csv results.

### RNA secondary structure

The in silico structural equivalence of the gecko and the human sequences was analyzed using CROSSalign (Ponti et al., 2018). First, the shorter profile of each couple was searched in the bigger one using the OBE-DTW procedure. Secondly, we performed Standard-DTW analysis to assess the structural alignment of the equivalent regions. The main output of this analysis is the structural distance, between the two input structures. The closer the distance is to 0 (with 0 meaning identical structural profiles) the higher the similarity in term of secondary structure. RNA molecules with a structural distance of 0.095-0.10 or higher are to be considered different in terms of secondary structure. For control comparisons in CROSSalign we generated a random lncRNA sequences of the same dinucleotide frequency (even the same number of each dinucleotide) as that of gecko RNAs using the web server of the Clote computational biology lab (http://clavius.bc.edu/~clotelab/RNAdinucleotideShuffle/). These random sequences were used to perform the Standar-DTW structural distance analysis with the human lncRNAs.

A Jupyter Notebook has been uploaded on GitHub to follow all the steps for k-mer and CROSSalign analysis in a Python3 environment: https://github.com/rodrisenovilla/Olazagoitia-Garmendia/blob/ffbeb12b889791b4b0c24f47cff4436cce2979eb/seekr.ipynb

For mobility assessment, the structural homologue regions from human and gecko *EVX1AS* were in vitro transcribed (IVT) using a T7 RNA Polymerase (Takara Bio, San Jose, CA). IVT lncRNAs were then purified, heated at 95ºC for 3 minutes, placed on iced and run in a non-denaturing agarose gel in TBE.

### Linear sequence similarity analysis

Dot plots were generated using EMBOSS dotmatcher (Rice et al., 2000). We used a window size of 5 and a threshold of 25.

mVISTA was used to visualize linear sequence alignments (Frazer et al., 2004). We used a 100bp sliding-window with a 25-bp nucleotide increment for the analysis of the underlying genomic alignment. Similarity was set at 70%.

### Synteny analysis (Genomic loci analysis)

Although current genome version annotation (Yamaguchi et al., 2021) is under construction, we have explored the synteny of the analogue lncRNAs of interest in this chromosome-scale genome version. Based on the v1 version, the gecko gene of interested was located by BLASTn (Altschul et al., 1990; Camacho et al., 2009) in the v2. We visualized the closest genes by Integrative Genome Viewer (IGV, https://igv.org/) (Robinson et al., 2011) and we retrieved the functional annotation of the closest genes by EggNOG () (Cantalapiedra et al., 2021). When no gene homologue was identified regarding the EggNOG standards, the default gene name given was kept (*e.g*. g13819).

### Protein-RNA binding motif analysis

The sequences of human and gecko *EVX1AS* were analyzed using MEME Motif Discovery tool in order to find enriched motifs within the lncRNAs. Additionally, TOMTOM Motif comparison tool was used to find proteins that would bind the enriched RNA motifs (Bailey et al., 2015).

### Subcellular localization

For quantification of *EVX1AS* levels in nuclear and cytoplasmic compartments, nuclei were isolated using C1 lysis buffer as described previously (Castellanos-Rubio et al., 2016) and amount of specific nuclear RNA measured by RT-QPCR was compared to the total amount of the RNA in the whole cell.

### Quantification of molecules per cell

In order to determine the *EVX1AS* copy number in human and gecko cells, a reference plasmid incorporating the cDNA sequence of each *EVX1AS* form was used. Absolute quantification was performed using 5 ten-fold serial dilutions of the reference standard. Ct versus the dilution factor was plotted in a base-10 semi-logarithmic graph, fitting the data to a straight line. Plot was then used as a standard curve for extrapolating the number of molecules of *EVX1AS* in the cells.

### Gecko tissue dissection

Three pre-hatching geckos at embryonic days (E)50 to E55 were anesthetized by hypothermia. After the opening of the egg, the embryos were sacrificed by cervical dislocation and a collection of tissues were obtained and flash frozen in liquid nitrogen: brain, heart, eyes, gut, lungs, liver, tail and the remaining carcase (containing mainly bones, muscle and skin).

### RNA tethering

CRISPR-display was performed as previously described (Shechner et al., 2015). Briefly, human and gecko lncRNAs fused to a U1 3’box at their 3’end and to a scaffold at their 5’ end were ordered as gBlocks (IDT). Subsequently, sgRNAs targeting the human or chicken *EVX1* TSS (available under request) were introduced by PCR and the whole construct was cloned into a pCMV plasmid. Then, the fusion lncRNA-sgRNA constructs were co-transfected with the catalytically inactive dCas9 into SHSY5Y cells or chicken embryos (see following section).

### Cells

The neuroepithelial SHSY5Y cell line (CRL-2266) was purchased from ATCC (Manassas, VA, US). Cells were cultured in 50% EMEM and 50% F12 medium supplemented with 10 % FBS (Millipore, Burlington, MA, USA #S0115), 100 units/ml penicillin and 100 μg/ml streptomycin (Lonza, #17-602E). For overexpression experiments 250 ng of each plasmid were used. 150000 cells/well were seeded and transfected using X-TremeGENE HP DNA transfection reagent (Sigma-Aldrich, #6366546001), cells were harvested after 48 h.

### Animals

All animal experiments were approved by a local ethical review committee and conducted in accordance with personal and project licenses in compliance with the current normative standards of the European Union (Directive 2010/63/EU) and the Spanish Government (Royal Decrees 1201/2005 and 53/2013, Law 32/107). Fertilized hen eggs (Gallus gallus), obtained from Granja Santa Isabel (Córdoba, Spain), were incubated at 37.5 °C in a humidified atmosphere until the required stages (Bellairs and Osmond, 2014). The day when eggs were incubated was considered embryonic day (E) 0.

Gecko eggs were harvested from a local breeder colony of Madagascar ground geckos (Paroedura pictus) at Achucarro (based on the colony at the Department of Ecology of Charles University, Czech Republic). Adult geckos were maintained on a 12/12-h light/dark and temperature cycle (8 a.m. lights on; 28 ° C diurnal temperature, 23 ° C nocturnal temperature) and provided with ad libitum access to food and water. Eggs were incubated at 28 ° C in a low-humidified atmosphere until the required stages (Noro et al., 2009). The day when eggs were found in the terrarium was considered E0.

### In ovo electroporation

Electroporation of chick embryos was performed as previously described (García-Moreno et al., 2014). Briefly, eggs were incubated in a vertical position at 38°C. Plasmids were injected with a volume of less than 1 μl into the fourth ventricle of E3 chick embryos using a fine pulled glass needle. Four electric pulses (14-17 V, 15 ms pulses with a 950 ms interval: BTX electroporator ECM) were then applied to the brain between insulated silver 40mm x 0.8mm wire electrodes with flattened pole (Intracel). Drops of Ringer’s solution supplemented with antibiotics (penicillin/streptamycin: Sigma) were added to the egg. Embryos were incubated until E6, when tissue was harvested for further research.

### RNA extraction and RT-QPCR

RNA from all samples was extracted using Direct-zol RNA miniprep kit (Zymo research, Irvine, USA, #R2053) with DNAse treatment. For the extraction of RNA from gecko and chicken tissues samples were homogenized with a pellet pestle prior to extraction.

500-1000 ng of RNA were used for the retrotranscription reaction using iScript cDNA Synthesis Kit (BioRad, CA, USA, #1708890). Expression values were determined by q-PCR using Sybr Green (iTaq SYBR Green Supermix, Bio-Rad, #1725124) and specific primers. *RPLP0* gene was used as endogenous control in human and in gecko and *Rplp7* in chicken samples. Reactions were run in a BioRad CFX384 and melting curves were analyzed to ensure the amplification of a single product. All qPCR measurements were performed in duplicate and expression levels were analyzed using the 2–∆∆Ct method. To reduce the variability across experiments, we normalized the relative expression as follows: all values from the same experiment were normalized to the highest value, hence we obtained values ranging from 0 to 1. In the case of *in vivo* experiments, relative expression values were normalized using z-score to allow the combination of values from different qPCRs. All primer sequences are listed in Supplementary Table 1.

### RNA immunoprecipitation (RIP)

For RIP experiments, SHSY5Y cells were lysed in RIP buffer (150 mM KCl, 25 mM Tris, 0.5 mM DTT, 0.5 % NP-40, PI), kept on ice for 15 minutes and homogenized using a syringe. IVT lncRNAs were incubated with RNA secondary structure buffer and added to the lysates. The mixes were pre-cleared with proteinG dynabeads (ThermoFisher, Waltman, MA, USA) for 1 h in a wheel shaker at 4ºC. Pre-cleared lysates were incubated with 1ug of MED1 antibody (Santa Cruz Biotechnologies, Dallas, TX, USA) for 1h at room temperature. After incubation dynabeads were added and further incubated for 30 min. The immunoprecipitation was washed three times with RIP buffer, three times with low salt buffer (50 mM NaCl, 10 mM Tris-HCl, 0.1 % NP-40) and three times with high salt buffer (500 mM NaCl, 10 mM Tris-HCl, 0.1 % NP-40). After the washes, 70 % of beads were resuspended in RNA extraction buffer and 30 % was used for WB.

### Western Blot

Laemmli buffer (62 mM Tris-HCl, 100 mM dithiothreitol (DTT), 10 % glycerol, 2 % SDS, 0.2 mg/ml bromophenol blue, 5 % 2-mercaptoethanol) was added to the protein samples and were denatured by heat. Proteins were migrated on 8% SDS-PAGE gels. Following electrophoresis, proteins were transferred onto nitrocellulose membranes using a Transblot-Turbo Transfer System (Biorad) and blocked in 5 % non-fatty milk diluted in TBST (20 mM Tris, 150 mM NaCl and 0.1 % Tween 20) at room temperature for 1 h. The membranes were incubated overnight at 4ºC with primary antibodies diluted 1:500 in TBST. Immunoreactive bands were revealed using the Clarity Max ECL Substrate (BioRad, #1705062) after incubation with a horseradish peroxidase-conjugated anti-mouse (1:10000 dilution in 2.5 % non-fatty milk) secondary antibody for 1 h at room temperature. The immunoreactive bands were detected using a Bio-Rad Molecular Imager ChemiDoc XRS and quantified using the ImageJ software (BioRad).

The following antibodies were used for Western Blotting: GAPDH (sc-47724) and MED1 (sc-74475).

### Immunohistochemistry

Embryonic chick brains were fixed by immersion in PFA (4% paraformaldehyde, PFA, diluted in phosphate buffered saline 0.1M– PBS, pH 7.3). Brains were transferred to PBS 6h after fixation. Brains were sectioned in the coronal plane at 50-70 μm thickness in a vibrating microtome (Leica VT1000S). Single and double immunohistochemical reactions were performed as described previously (Rueda-Alanã and Garciá-Moreno, 2021) using the following primary antibodies: rabbit antibody to histone H3 - phospho S10 (Abcam ab47297; 1:1000), mouse antibody to LHX5 (DSHB;1:30) and mouse antibody to Meis2 (DSHB; 1:30). For secondary antibodies (all 1:1000), we used Alexa 647 goat antibody to rabbit IgG (Molecular Probes, A32733) and Alexa 488 goat antibody to mouse IgG (Molecular Probes, A11001). Sections were counterstained with DAPI.

### Statistical analyses

The data are represented as the mean ± standard error of the mean of at least three biological replicates. Mean comparisons were performed by Student’s t-test. The statistically significance level was set at p< 0.1.

## Code availability

Although many of the analysis were carried out in web-page apps, we have coded a pipeline to unify all our bioinformatic analysis (<u>Olazagoitia_pipeline.ipynb</u>) and to generate the boxplots for SEEKR results (<u>seekr_output_explore.Rmd</u>). These files can be found in our GitHub (https://github.com/rodrisenovilla/Olazagoitia-Garmendia). If you would like further explanations or find trouble re-running them, contact us for support.

## Acknowledgements

This work was supported by grants PGC2018-097573-A-I00 from the Spanish Ministry and COLLAB20/02 from the UPV-EHU to ACR; PGC2018-096173-A-I00 grant from the Spanish Ministry and PIBA 2020_1_0057 grant from the Basque Government to FGM.

AOG was supported by a predoctoral fellowship from the Basque Government. RSG is supported by a predoctoral fellowship by Fundación Tatiana Pérez de Guzmán el Bueno.

## SUPPLEMENTARY MATERIALS

### Supplementary Figures

**Figure S1:**
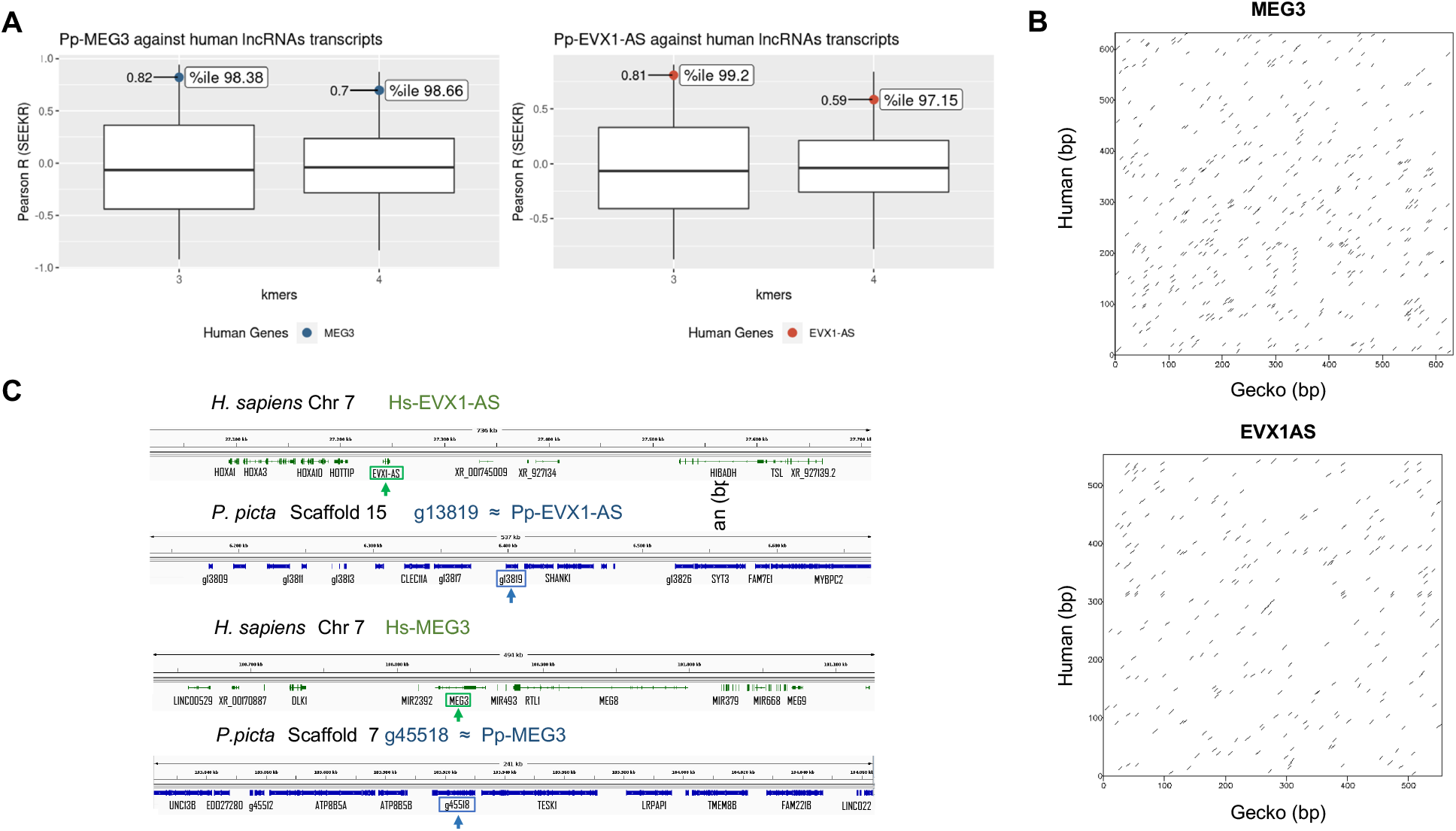
**(A)** k=3 and k=4 k-mer content analysis of *Pp-MEG3* and *Pp-EVX1AS* candidates. Candidate lncRNAs were compared against the human lncRNA repository (Gencode v1). Blue and red dots correspond to the *Pp-MEG3* and *Pp-EVX1AS* genes respectively. **(B)** Linear similarity of the human and gecko *MEG3* and *EVX1As* lncRNAs by Dot Plot analysis. **(C)** Synteny analysis of gecko and human. Top: Genomic locus for *EVX1-AS* human gene (green) and locus of its functionally equivalent gecko’s gene, g13819 ≈ *pp-EVX1-AS* (blue). Bottom: Genomic locus for *MEG3* human gene (green) and locus of its functionally equivalent gecko’s gene, g45518 ≈ *pp-MEG3* (blue). Genome visualization is carried out by Integrative Genome Viewer (IGV) with Human (GRCh38/hg38, RefSeq.gtf) and P. picta (v2, BRAKER.gtf) genomes, respectively.

**Figure S2:**
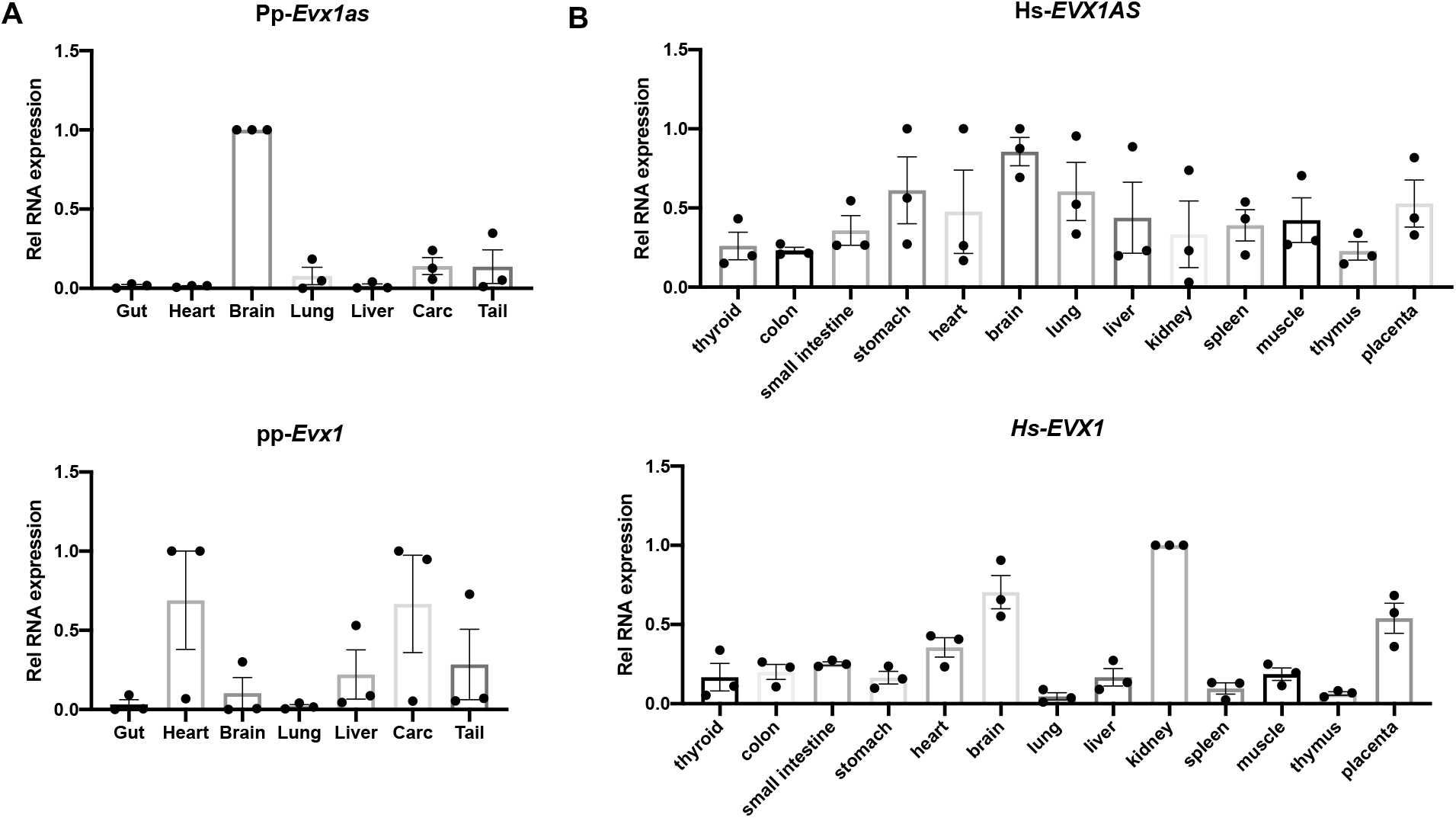
Relative expression of **(A)** gecko *Meg3, Evx1as* and *Evx1* and **(B)** human *EVX1AS* and *EVX1* in a range of tissues. Data represents the mean and standard error of three independent experiments.

**Figure S3:**
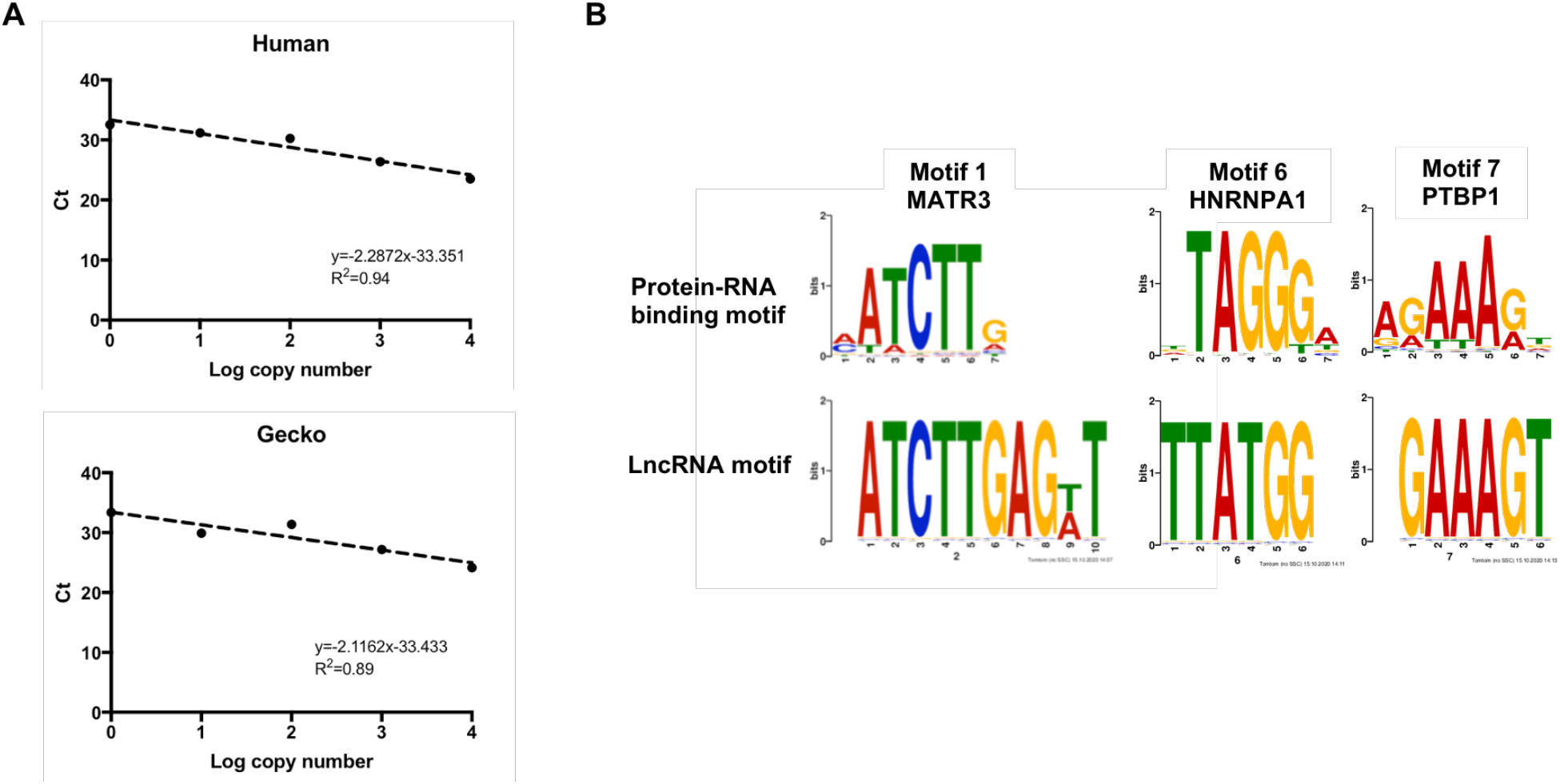
**(A)** Standard curve for the human (top) and gecko (bottom) *EVX1AS* copy number analyses. **(B)** Alignment of the enriched lncRNA motifs with described RNA binding protein motifs using TOMTOM Motif comparison tool.

**Figure S4:**
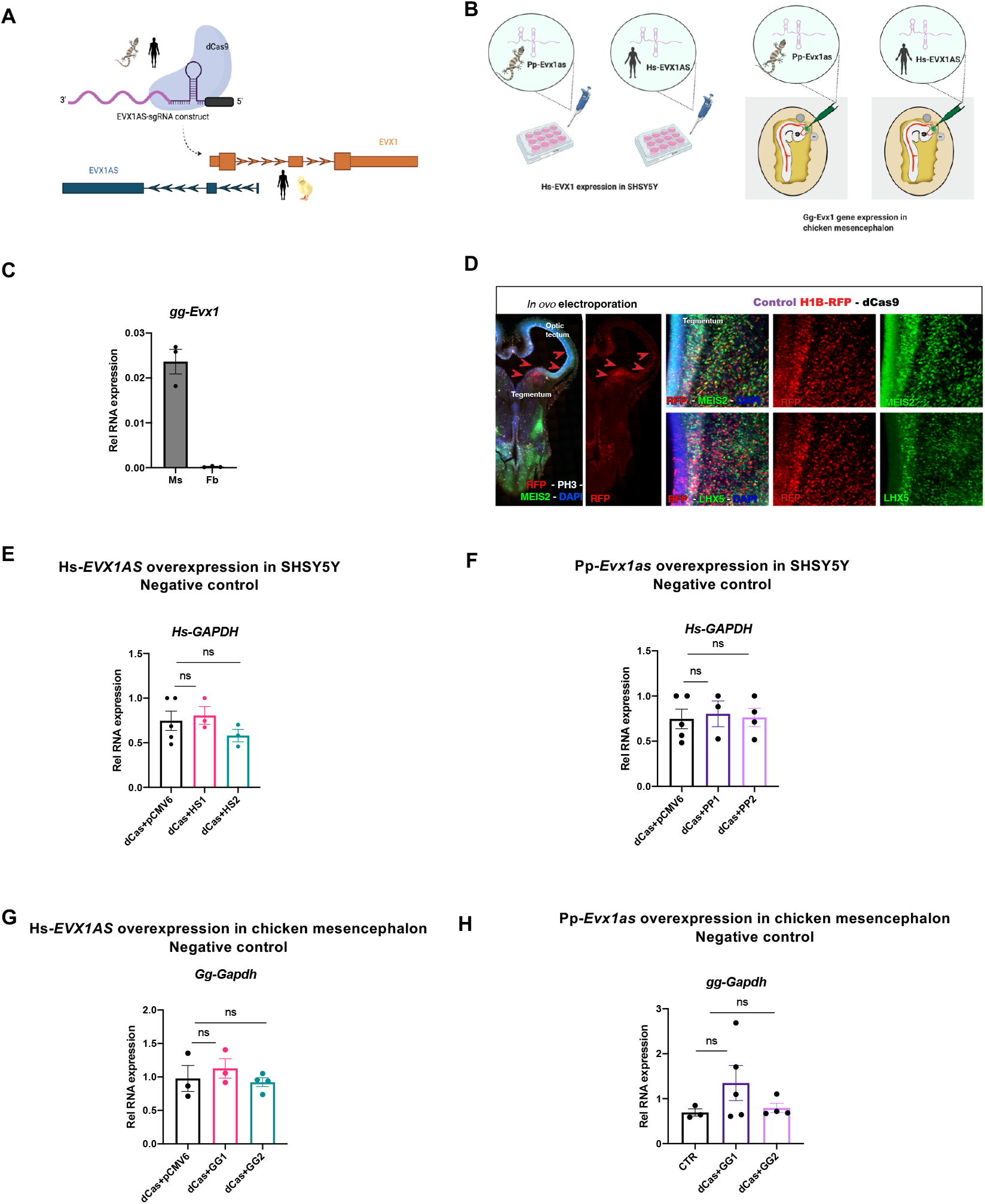
**(A)** Schematic representation of the RNA tethering experiments. **(B)** Schematic representation of the *in vitro* (left) and *in vivo* (right) experiments. **(C)** *Evx1* expression analysis in chicken embryonic mesencephalon (Ms) and forebrain (Fb). Data represents the mean and standard error of brain tissue from 3 chicken embryos. **(D)** Immunohistochemistry of the chicken embryonic midbrain on coronal section showing the successful transfection of the control plasmid. Transfected cells are shown in red due to their ectopic expression of red fluorescent protein RFP. LHX5 and MEIS2 patterns of expression (in green) indicate the mesencephalic region that was actually transfected. PH3 immunostaining shows mitotic cells at the germinative zone of the mesencephalon, in white. DAPI counterstain in blue. Relative expression of *GAPDH* negative control in SHSY5Y cells after *Hs-EVX1AS* **(E)** and *Pp-Evx1as* **(F)** overexpression and in chicken mesencephalon after *Hs-EVX1AS* **(G)** and *Pp-Evx1as* **(H)** overexpression. Data represents the mean and standard error of at least three independent experiments. ns: non significant. HS1, HS2 and GG1, GG2 correspond to the two different sgRNAs used for human and chicken transfections respectively.

### Supplementary Tables

**Supplementary Table 1.**
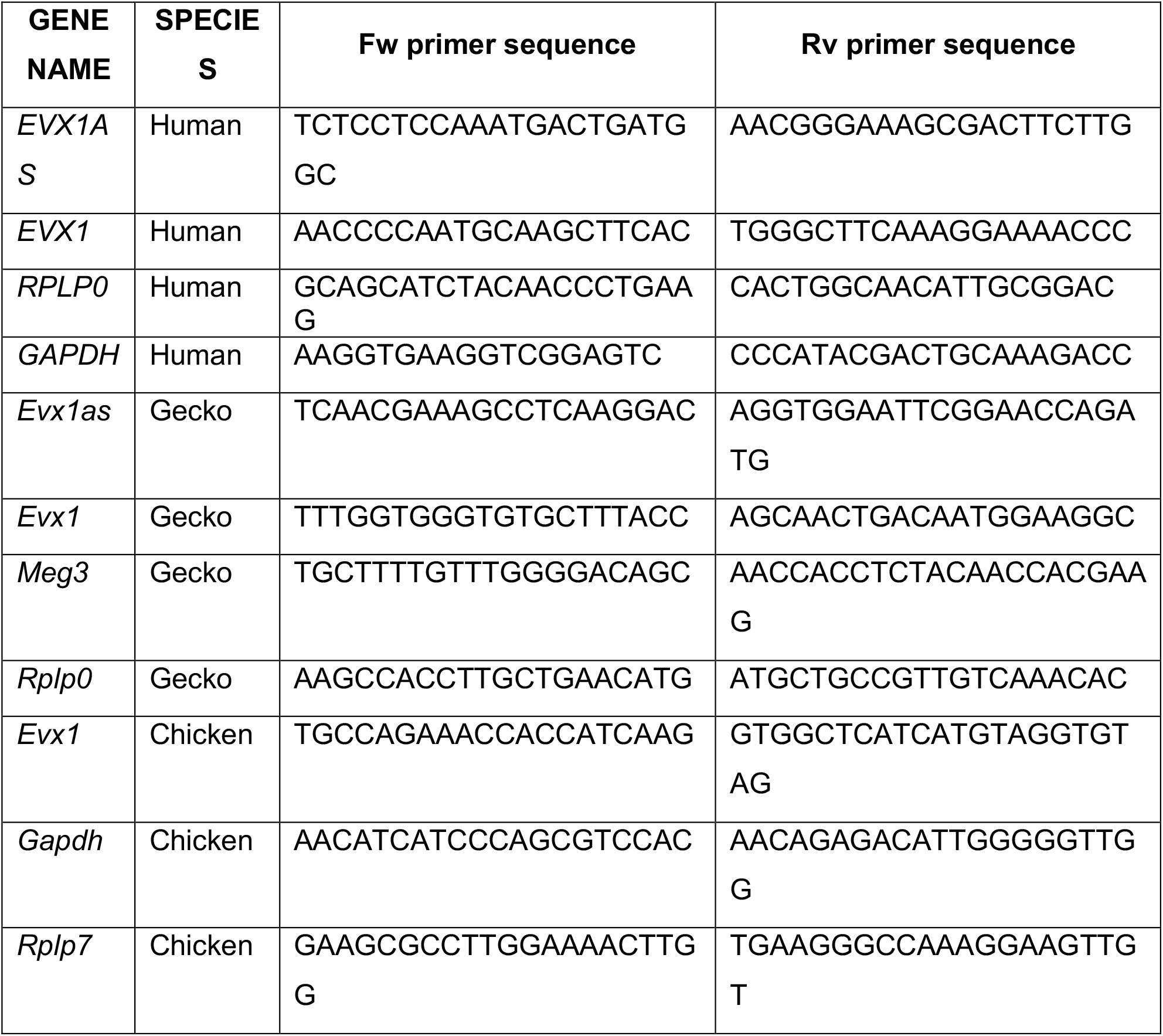
List of used primers and sequences.

## Notes

### Competing Interest Statement

The authors have declared no competing interest.

### Summary of Updates

In this version of the manuscript, we have performed new analyses and added new figures that better show the distribution of the k-mer content for the candidate gecko lncRNAs. We have now included more details into the methods sections and we have generated a Jupyter notebook that we have uploaded to Github with all the steps of our in silico analyses. We have also added more description in the results, together with the addition of graphs showing independent data points. Note that the actual list of authors has changed and that we have included Rodrigo Senovilla-Ganzo as a co-first author, as he has been a key piece for all the new in silico analyses that we have included in the revised version of the manuscript.

